# Simultaneous Determination of the Size and Shape of Single α-Synuclein Oligomers in Solution

**DOI:** 10.1101/2023.01.09.523202

**Authors:** Saurabh Awasthi, Cuifeng Ying, Jiali Li, Michael Mayer

**Author notes:** Corresponding Author Email ID.

## Abstract

Soluble oligomers of amyloid-forming proteins are implicated as toxic species in the context of several neurodegenerative diseases. Since the size and shape of these oligomers influences their toxicity, their biophysical characterization is essential for a better understanding of the structure-toxicity relationship. Amyloid oligomers are difficult to characterize by conventional approaches due to their heterogeneity in size and shape, their dynamic aggregation process, and their low abundance. This paper demonstrates that resistive-pulse measurements using polymer-coated solid-state nanopores enable single-particle level characterization of the size and shape of individual αSyn oligomers in solution within minutes. A comparison of the resulting size distribution with single-particle analysis by transmission electron microscopy and mass photometry reveals that nanopore-based characterization agrees well with both methods, while providing better size resolution and elucidating that αSyn samples are composed of stable oligomer sub-populations that contain multiples of approximately 12 monomers (i.e., 12-, 24-, 48-, 60-, 84-mers). Applying the unique capability of nanopores to approximate particle size and shape to picomolar concentrations of αSyn oligomers in the putatively toxic size range, revealed shapes that agree well with previous estimates by cryo-EM with the added advantage that nanopore-based analysis occurs rapidly, in solution, and has the potential to become a widely accessible technique.

## Introduction

Neurodegenerative diseases characterized by misfolding and aggregation of α-synuclein (αSyn) protein (14.5 kDa) are collectively termed synucleinopathies.^1–4^ The most common types of synucleinopathies are Parkinson’s disease (PD), dementia with Lewy bodies (DLB), and multiple system atrophy (MSA).^3^ Oligomers of αSyn are neurotoxic species and implicated in the pathology of Parkinson’s disease.^5–9^ Colla et al. showed that αSyn accumulates within the endoplasmic reticulum or in microsomes forming toxic oligomers in transgenic mouse brain leading to α-synucleinopathy.^10^ Prots *et al*. demonstrated that αSyn oligomers disrupt axonal integrity in human neurons and that increased αSyn oligomerization by expressing oligomer-forming mutants (E46K and E57K) of αSyn resulted in impaired axonal transport of mitochondria.^11^

Although most soluble oligomeric species of αSyn are toxic,^9^ the molecular basis of toxicity remains to be established.^12^ Therefore, the biophysical characterization of αSyn oligomers is essential for understanding their pathology and developing therapeutic interventions. Characterization and quantification of αSyn oligomers, remains however, challenging with established approaches due to the heterogeneity of oligomeric species with regard to their size, shape and their low abundance with concentrations ranging from picomolar (pM; 10^-12^ mol l^-1^) to nanomolar (nM; 10^-9^ mol l^-1^).^13, 14^ One reported mechanism of toxicity of αSyn oligomers involves interactions with lipids and formation of pores in phospholipid membranes.^15–17^ The physical properties of αSyn oligomers such as oligomer size or conformation/shape are key determinants of their pore-forming activity and toxicity.^17–22^ Specific sizes (i.e., oligomers with a certain number of monomers) or specific shapes (such as annular or disc-shaped) oligomers have been associated with neurotoxicity in vivo. ^9, 17, 20^ Recently, Kiechle et al. used in vivo protein complementation to identify presynaptic oligomerization and age-dependent accumulation of potentially toxic αSyn oligomers in a specific size range (8-16-mer).^23^ Also, prior studies have identified and demonstrated neurotoxic αSyn oligomers composed of ~30 monomers,^24^ >30 monomers and 10-40 monomers.^20, 25^

Characterization methods based on ensemble analysis, such as size exclusion chromatography (SEC), analytical ultracentrifugation (AUC), small-angle X-ray scattering (SAXS), dynamic light scattering (DLS), and gel electrophoresis, fail to capture the entire range of heterogeneity in size with good resolution and they either cannot determine oligomer shape or struggle to do so in heterogenous mixtures. In addition, the need for fluorophore labeling, cross-linking or preparation of dry samples of some existing methods may result in undesirable alterations in the physical and biochemical properties of amyloid oligomers.^26, 27^ Recently, Arter et al. used microfluidic free-flow electrophoresis as a solution-based approach to fractionate stable αSyn oligomer ensembles for achieving structural, kinetic and immunological characterization in addition to the measurement of average oligomer ζ-potential of each fraction.^28^ Alternative, attempts have employed fluorescence methods such as single-molecule Förster resonance energy transfer (smFRET),^21^ and single-molecule photobleaching to characterize αSyn oligomers in vitro with respect to oligomer size.^29^ The goal of the work presented here was to address the challenge of analysing heterogenous oligomer samples by establishing an accessible and practical method for characterizing αSyn oligomers on a single oligomer level, rapidly, and in solution.

An elevated level of αSyn oligomers in cerebrospinal fluid (CSF) or plasma is associated with the onset and progression of neurodegenerative synucleinopathies,^30, 31^ therefore, αSyn oligomers are promising biomarker candidates.^30, 32^ Commonly used immunoassays to quantify amyloid oligomers such as enzyme-linked immunosorbent assay (ELISA),^33, 34^ single-molecule array (SIMOA) assays^35, 36^ and surface-based fluorescence intensity distribution analysis (sFIDA)^37^ require the use of high-affinity antibodies and typically fail to discriminate oligomers from monomers and fibrils. Moreover, a recent study by Kumar et al. tested sixteen so-called “conformation-specific αSyn antibodies,” and none of these antibodies showed specificity for any particular aggregate size of αSyn.^35^ Given the current challenges associated with immunological assays for oligomer quantification, an urgent need exists for an antibody-free, sensitive approach that can determine oligomer size, abundance and ideally shape.

We propose that one attractive approach to address this need is to monitor aggregation using nanopores.^38–46^ So far, most studies using nanopores focused on either the detection of particles or on monitoring the aggregation process over time, typically these studies did not determine the size of amyloid oligomers and even fever studies interrogated oligomer shapes.^39, 41, 46–48^ For a review of progress in this field, please see Houghtaling et al^46^ as well as recent reports.^40, 41, 43, 44, 49–53^ Here, we apply this approach to single-particle characterization of oligomers of αSyn and reveal their size and shape. We demonstrate that the size distribution of αSyn oligomers contains ten sub-populations of oligomers with distinct and stable number of aggregated monomers. Supported by TEM-based single-particle imaging and mass photometry analysis, these stable oligomer states start with dimers, trimers or tetramers, and then appear to contain multiples of approximately 12 monomers including 12-mers, 24-mers, 48-mers 60-mers and 84-mers. Finally, we demonstrate that the size and shape analysis of individual oligomers makes it possible to detect and quantify oligomers of specific size, which have been identified as “toxic oligomers” such as 10S and 15S^20^ and to reveal the distribution of the shape of these oligomers. In this way, the abundance of a sub-population with putatively toxic size and putatively toxic shape (such as prolate-shaped or tubular oligomers^17, 20^ that possess pore forming activity) may be determined.

## Results and Discussion

### Analysis of the size of α-synuclein oligomers at the single-particle level

**Figure 1A** illustrates the experimental setup for resistive pulse measurements using solid-state nanopores. A single pore with a diameter of 26 nm to 56 nm in a thin insulating silicon nitride (SiN_x_) membrane separates two electrolyte compartments. A high-gain, low-noise patch-clamp amplifier applies a potential difference (±0.1 V) between these compartments and monitors the ionic current through the nanopore. Passage of individual αSyn oligomers through the nanopore displaces conducting electrolyte, resulting in characteristic resistive pulses. These resistive pulses contain information about the physical properties of the translocating particles, including their volume (i.e., size ‘*Λ*’) and shape (i.e., length-to-diameter ratio ‘*m*’ of a corresponding ellipsoidal model) (**see Figure 1B**).^50^ The dwell time that a protein resides in the pore, *t_d_*, is influenced by its electrophoretic mobility and hence its net charge. In addition, the frequency of observed translocation events is proportional to the concentration of oligomers.^54^

**Figure 1.**
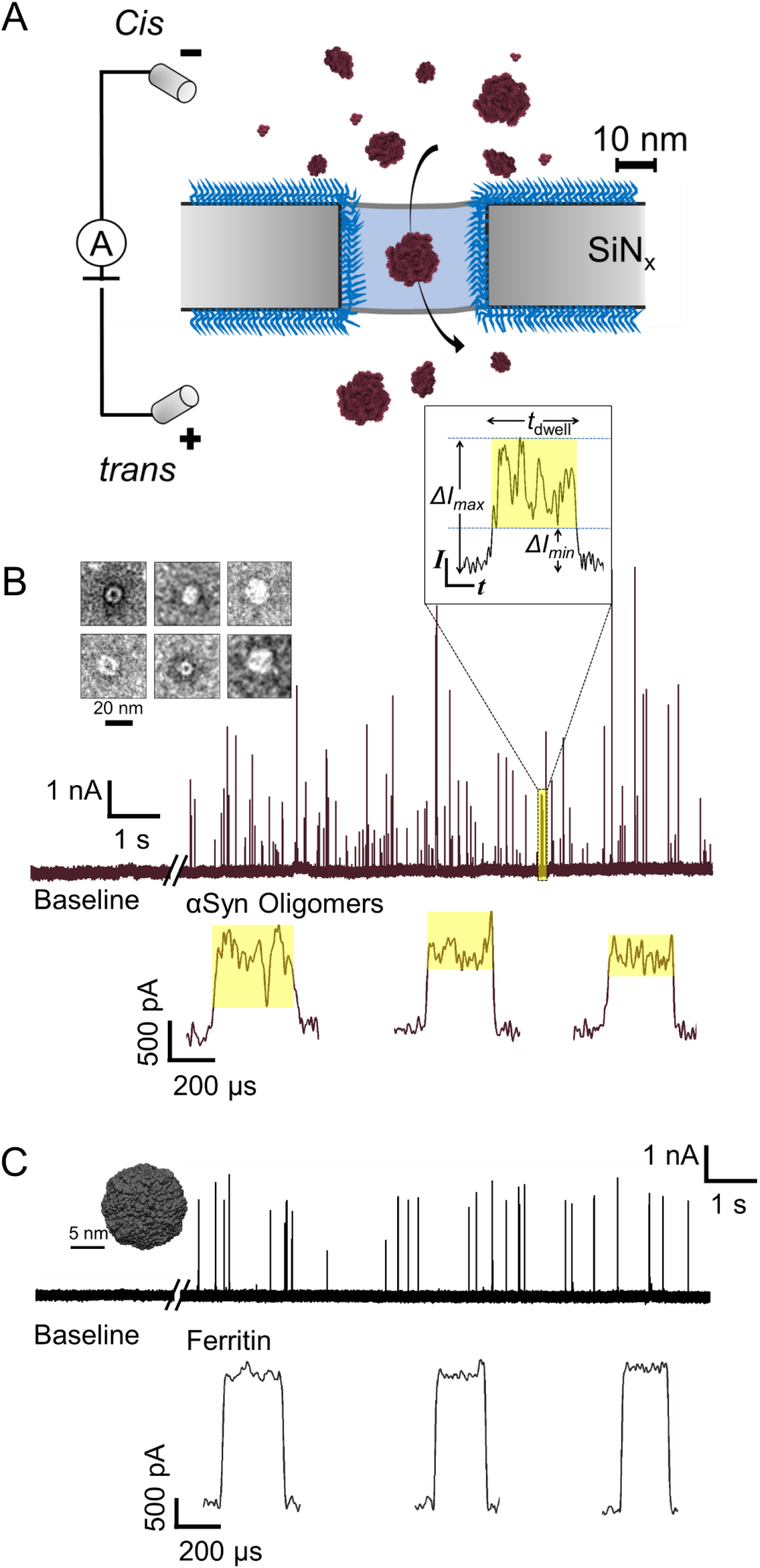
Nanopore-based, single-particle characterization of α-synuclein oligomers in solution. **A**. Schematic illustration of the experimental setup to measure resistive pulses due to the translocation of individual αSyn oligomers through a polymer-coated nanopore (blue). **B.** Original current recordings before and after adding αSyn oligomers show resistive pulses due to the translocation of αSyn oligomers as upward spikes. The TEM image in the inset shows individual αSyn oligomers from the sample used for nanopore experiments. Characteristic translocation events from the passage of individual αSyn oligomers through the nanopore illustrate the dwell time (*t_d_*), minimum current blockade (*Δ*/_min_), and maximum current blockade (*Δ*/_max_) that arise from the rotational dynamics of non-spherical oligomers translocating through the electric field inside the nanopore. The magnitude of current blockade *Δ*/ is proportional to the particle volume, while the intra-event current modulations (shaded in yellow) contain information about the shape (*m*) and orientation of each oligomer in solution.^50, 51^ Three additional examples of translocation events (*t_d_* >150 μs) from the passage of individual αSyn oligomers through a nanopore are shown below. **C**. Original current recordings before and after adding the perfectly spherical protein ferritin (PDB ID: 6TSF) show resistive pulses as upward spikes due to single protein translocations. Note that the three examples of translocation events that are shown in detail (*t_d_* >150 μs) from the passage of ferritin through the same nanopore result in square-shaped resistive pulses with small intra-event current modulations. For this work, we used nanopores that resulted in such square-shaped resistive pulses for ferritin translocations, indicating that these pores correctly detect small current modulations from rotations of a perfectly spherical protein. In this case the modulations are similar in amplitude to the baseline noise as reported before.^50, 51^

**Figure 1B** shows original current traces of electrical recordings through nanopores, featuring translocation events of individual oligomers as resistive upward spikes (i.e. reductions in the amplitude of the negative baseline current). In order to evaluate the benefits of nanopore-based oligomer characterization on a single particle level, we employed samples of αSyn oligomers obtained from ND Biosciences, Switzerland, which were prepared as explained by Kumar et al in order to be stable over time with regard to the size distribution of oligomers.^55^ To determine the size distribution of these samples, we recorded thousands of individual translocation events of αSyn oligomers. We classified αSyn oligomers based on their size in terms of the number of monomers they contain. To this end, we used the volume of monomeric αSyn protein, which we estimated to be 35 nm^3^ using nanopore experiments (**See Supplementary Note 1 and Supplementary Figure S1**), in good agreement with earlier reports of αSyn monomer volume (~30 nm^3^) determined by SAXS.^56^ Oligomer sizes estimated by nanopores range from a dimer to large-sized oligomers consisting of ~150 monomers (**Figure 2**), again in good agreement with the sizes of αSyn oligomers reported earlier.^20, 21, 57^ **Figure 2A** shows that single-oligomer analysis of these samples using nanopores, revealed 10 maxima in the size distribution corresponding to 10 different sizes within the oligomer population. We labeled different-sized sub-populations of oligomers as O^1^ to O^10^ with increasing size. Established single-particle methods, including negative staining TEM imaging (**Figure 2B**) and mass photometry (**Figure 2C**) confirmed the oligomer size estimates from the nanopore experiments. Mass photometry is a recently introduced method based on light-scattering that enables oligomer mass analysis at the single-particle level during their interaction with a surface.^58, 59^ **Figure 2** demonstrated that the size range of oligomer population determined by TEM (21- to 133-mer) and mass photometry (3- to 103-mer) agrees well with the range established by nanopore experiments (dimer to 122-mer). Nanopore-based analysis revealed two small-sized oligomer populations, called O^1^ and O^2^, corresponding to (4±3)-mer and (11±3)-mer αSyn oligomers, which were also detected by mass photometry but not by TEM imaging since they were too small (**Figure 2**). Comparing oligomer size estimations using nanopores to those using TEM and mass photometry shows the agreement between the range of estimated sizes from all three methods and illustrates that the resolution for distinguishing between differently-sized oligomers is highest from nanopore experiments, which reveals 10 sub-populations in size, while the resolution of TEM imaging and mass photometry are limited to approximately half the number of sub-populations.

**Figure. 2.**
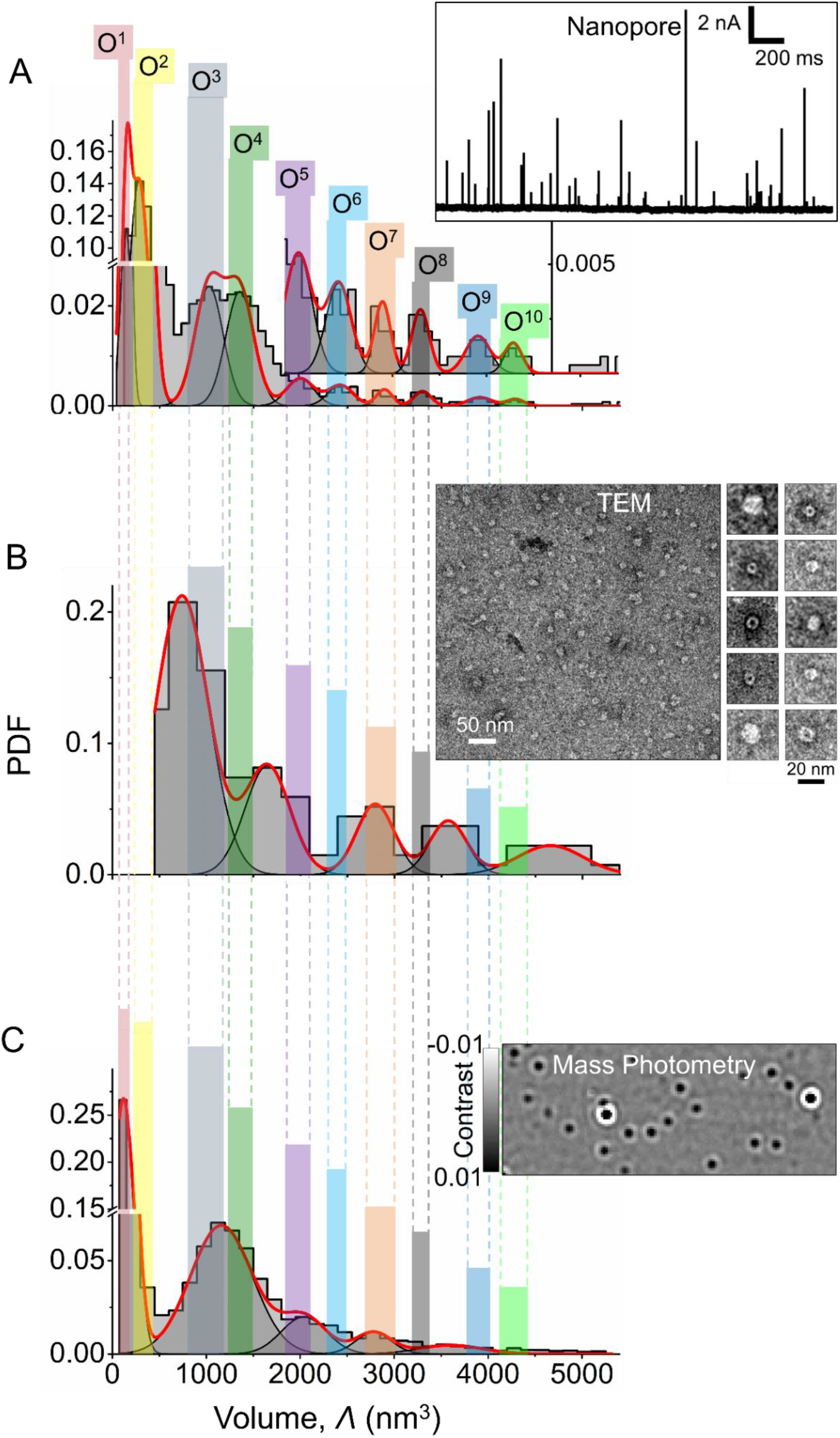
Nanopores enable high-resolution analysis of oligomer size. **A.** Size distribution of αSyn oligomers as determined by nanopores. Differently-sized oligomer sub-populations identified using the nanopore-based approach are marked as O^1^ to O^10^ and range in size from a (4±3)-mer to (122±5)-mer. This size distribution is the composite result from several nanopore experiments with different nanopore chips that had all been calibrated for accurate volume detection by translocating ferritin proteins. The inset shows original current recordings through a nanopore with resistive pulses due to the translocation of single αSyn oligomers as upward spikes. **B.** Size distribution of αSyn oligomers determined by TEM. The inset shows a TEM micrograph of αSyn oligomers used for the analysis. The resolution is limited because only a two-dimensional image can be obtained and the particle volume was estimated considering the long axis as the particle diameter of the 2D projection **C.** Size distribution of αSyn oligomers determined by single-particle mass analysis using mass photometry. The inset shows a differential interferometric scattering image of αSyn oligomers. For comparison, the shaded regions in different colors show the presence/absence of different oligomer populations identified by nanopores, TEM, and mass photometry. Since mass photometry is calibrated to reveal the molecular weight of each particle, we converted the molecular weight to particle volume using the molecular weight of αSyn monomers and assuming a monomer volume of 35 nm^3^.

In addition, we compared the size distribution of αSyn oligomers obtained by nanopore recordings with previously reported sizes of αSyn oligomers. One of the major oligomer populations identified by nanopores labeled as O^2^ consisting of 2-14 monomers, agrees well with αSyn oligomers containing 8-16 monomers recently reported to accumulate in the brain of transgenic mouse models with increasing age as determined by a protein complementation approach.^23^ Two other major αSyn oligomer populations labeled as O^3^ and O^4^ consisting of ~30 and ~39 monomers, agree well with earlier reports of ~30-mer αSyn oligomers as determined by size exclusion chromatography (SEC) coupled with multiangle laser light scattering (MALLS),^24^ SAXS,^60^ analytical ultracentrifugation (AUC),^61^ and single-molecule photobleaching experiments.^29^ We confirmed the presence of these αSyn oligomer populations by particle size analysis using TEM and mass photometry and highlighted them with light blue and light green shaded regions in **Figure 2B, 2C**. While mass photometry, like nanopore experiments, made it possible to identify small oligomers such as tetramers, its resolution was limited to five different oligomer sizes. We also used TEM for particle size analysis to determine oligomer size distributions but it could not detect the smallest oligomers (such as tetramers and octamers). Overall, the peaks in the size distribution from all three methods indicate that certain sizes of αSyn oligomers are more probable in the sample than others. The most prominent among these are oligomers composed of 4±3 monomers (O^1^, **Figure 2A, C**), 11±3 monomers (O^2^, **Figure 2A, C**), 25±10 monomers (O^3^, **Figure 2A, B, C**), 44±4 monomers (O^4^, **Figure 2 A, B, C**), 57±8 monomers (O^5^, **Figure 2A, B, C**) and 82±5 monomers (O^7^, **Figure 2 A, B, C**). These sizes suggest that the most stable αSyn oligomers are composed of multiples of 12±4 monomers, including 12-mers, 24-mers, 48-mers, 60-mers, and 84-mers.

### Analysis of the shape of individual αSyn oligomers

Nanopore-based resistive pulse recording makes it possible to approximate the spheroidal shapes of individual particles by analysing the current modulations during individual translocation events of single oligomers through the nanopore. ^50^ This approximation models particle shape with an ellipsoid with axes *A, B, B* of the same volume and returns the ratio, *m=A/B* between the two axes of the ellipsoid that best represents each individual αSyn oligomer.^51^ To this end, nanopore-based shape analysis exploits the minimum and maximum current blockades (Δ/_min_ and Δ/_max_ values, **Figure. 1B**), which correspond to the two extreme orientations of a non-spherical particle with certain volume and shape in the electric field of the nanopore.^50, 51^ Earlier studies examining the morphology of αSyn oligomers based on TEM, atomic force microscopy (AFM), and small angle X-ray scattering (SAXS) have revealed oligomers with spherical,^62, 63^ annular, toroidal, flattened disc-like, globular and prolate shapes.^20, 24, 56, 64, 65^ Here, based on the Δ/_min_ and Δ/_max_ values of each translocation event, we model the shape of each oligomer in one attempt as an oblate ellipsoid and in the other attempt as a prolate ellipsoid since solutions for both models can be found for many particles. TEM imaging of the same samples revealed the absence of any large fibrillar or elongated aggregates that may exceed the length of the nanopore (**Figure 2B**), making it possible to use the same analysis of volume-exclusion in the nanopore for characterizing all translocation events.^50, 51^ In addition, we confirmed that nanopore recording at the high ionic strength (i.e., 2 M KCl) that we used, does not change the size distribution of αSyn oligomers during the experiment compared to solutions with physiologic ionic strength (**Supplementary Note 2,** and **Supplementary Figure S2 and S3**).

Nanopore-based simultaneous determination of oligomer size and shape of each particle provides the opportunity to characterize oligomer sub-populations with respect to these two important parameters. For instance, we determined the shape of αSyn oligomers in a size range that was previously described to include toxic oligomers (**See Figure 3A-D**).^20, 23, 56^ Giehm et al. used SAXS experiments to determine a wreath shape with and average length-to-diameter ratio, *m* of 0.25 of so called “vesicle disrupting α-synuclein oligomers” consisting of ~16 monomers.^56^ **Figure 3A** shows the distribution of *m* values obtained for the prolate model of 16±1-mer oligomers by nanopore experiments and reveals that the major oligomer sub-population corresponds to an oblate shape value of *m* = 0.24 in excellent agreement with the reported shape of 0.25 from SAXS experiments.^56^ In addition to this dominant sub-population, nanopore-recordings identified a second, less prominent sub-population with a shape value of *m* = 0.64 in the same sample. Due to its low abundance, this sub-population may not be detectable in SAXS-based ensemble analysis.

**Figure. 3.**
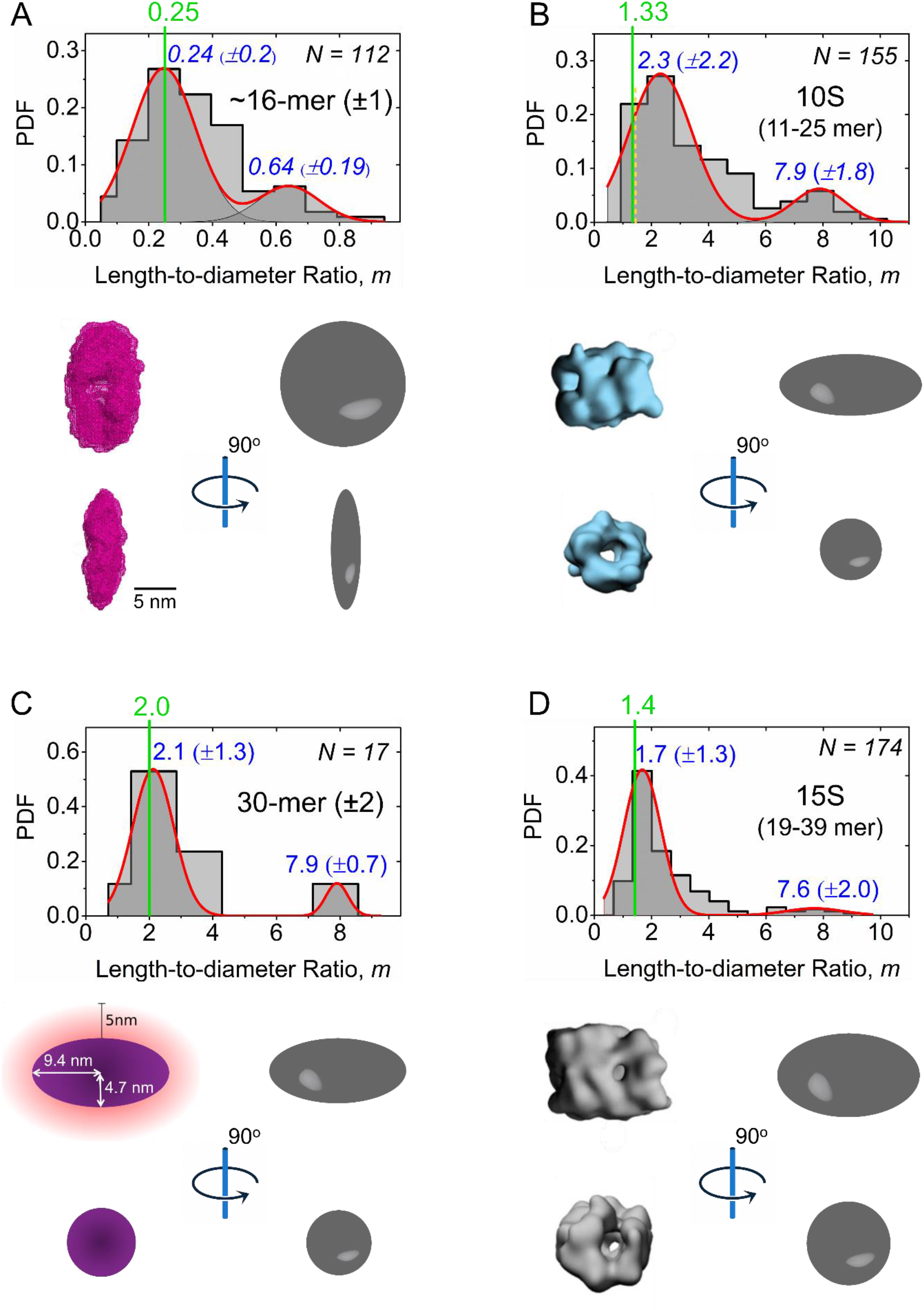
Shape approximation of putatively toxic αSyn oligomers in solution. **A**. Distribution of the length-to-diameter ratio, *m* of oligomers consisting of 15-17 αSyn monomers. The cartoon below shows a comparison of the approximated ellipsoidal shape of a ~16-mer determined by nanopore (grey ellipsoid) with the structure of the 16-mer revealed by SAXS experiments (mesh representation, in pink), adopted with permission from reference ^56^ **B**. Distribution of the length-to-diameter ratio of 10S oligomers consisting of 11-25 αSyn monomers. The cartoon below shows a comparison of the approximated ellipsoidal shape as determined by nanopore (grey spheroids) with the three-dimensional structures for 10S oligomers determined by cryo-EM, reprinted with permission from reference ^20^. The dashed line in yellow marks the first bin of the histogram, which represents the second most probable bin of the shape distribution for this oligomer size range (11-25 monomers) as determined by nanopore experiments and agrees well with the previously reported shape value (green line).^20^ **C**. Distribution of the length-to-diameter ratio of 30±2-mer oligomers. The cartoon below shows a comparison of the approximate ellipsoidal shape of 30-mers as determined by nanopore (grey spheroids) with an ellipsoidal model of the 30-mer determined by SAXS, reprinted with permission from reference ^24^ **D**. Distribution of the length-to-diameter ratio of 15S^20^ oligomers consisting of 19-39 αSyn monomers. Comparison of the approximate ellipsoidal shape as determined by nanopore (grey spheroids) with the three-dimensional structures for 15S oligomers determined by cryo-EM, reprinted with permission from reference ^20^. The most probable *m* value of each sub-population is shown at the top in blue. The solid lines in light green show the reference length-to-diameter ratio, *m*, for the respective oligomeric species.

**Figure 3B** and **3D** show the distribution of the approximated shape of so-called “kinetically trapped toxic αSynuclein oligomers” with two sizes: 10S (11-25 mer) and 15S (19-39 mer), which has previously been characterized as prolates with average values of *m* = 1.33 for 10 S oligomers and *m* = 1.4 for 15S oligomers based on their sedimentation coefficients.^20^ The nanopore experiments reveal a dominant oligomer sub-population with a length-to-diameter ratio, *m* of 2.3 for the 10S oligomers and a dominant oligomer sub-population with a length-to-diameter ratio, *m* of 1.7 for the 15S oligomer in good agreement with the previously reported average shape values of 1.33 and 1.4, respectively.^20^

In addition to these studies, Lorenzen et al. used size exclusion chromatography in combination with multi-angle scattering and SAXS and determined an average prolate ellipsoid shape with a *m* value of 2.0 for oligomers consisting of 30±2 monomers.^24^ The nanopore experiments carried out here, indicate a major sub-population with a *m* value of 2.1 in the distribution of the length-to-diameter ratio of 30-mer oligomers (**Figure 3C**), again in excellent agreement with the reported average shape value. Overall, **Figure 3** indicates that for oligomeric species within size brackets that have previously been reported as toxic, nanopore recordings resolve typically one dominant sub-population in oligomer shape that agrees well with the previous reports as well as a less prominent sub-population of oligomer shape that might be missed using ensemble analysis.

### Quantifying αSyn oligomers

In order to assess if nanopore-based analysis is able to quantify the total concentrations of αSyn oligomers as well as the concentration of sub-populations, we monitored the frequency of individual translocation events through nanopores at increasing total oligomer concentrations ranging from picomolar (pM; 10^-12^ mol l^-1^) to nanomolar (nM; 10^-9^ mol l^-1^). **Figure 4A** shows that the frequency of translocation events increased linearly with increasing total oligomer concentration. **Figure 4B** shows that the translocation event frequency is directly proportional to the total concentration of αSyn oligomers at least up to ~2.5 nM (**See Figure 4B**). This sensitivity is beneficial for assessment of the concentration of αSyn oligomers as a potential biomarker for PD. We propose that PD biomarker analysis will benefit from the ability to count individual oligomer translocations using the nanopore-based method because it is not limited to quantifying the total concentration of oligomers but also makes it possible to identify and quantify the fraction of sub-populations. For instance, it can determine the concentration of αSyn oligomers of certain sizes such as 8-10 mers, 21-39 mers, 40-50 mers, which are considered toxic. Or it can quantify the fraction of oligomers with a certain shape or the combination of a certain size and shape. **Figure 4B** shows, for instance, that the frequency of translocations from the specific-sized oligomers consisting of 29±5-monomers or from oligomers with a shape value ranging from 1.4 to 2.0 also increased linearly with total oligomer total oligomer concentration. This capability of quantifying specific sub-populations is a direct consequence of the characterization of each individual translocation event with respect to the size and shape of the oligomer particle that caused the resistive pulse combined with its measured frequency of occurrence.

**Figure. 4.**
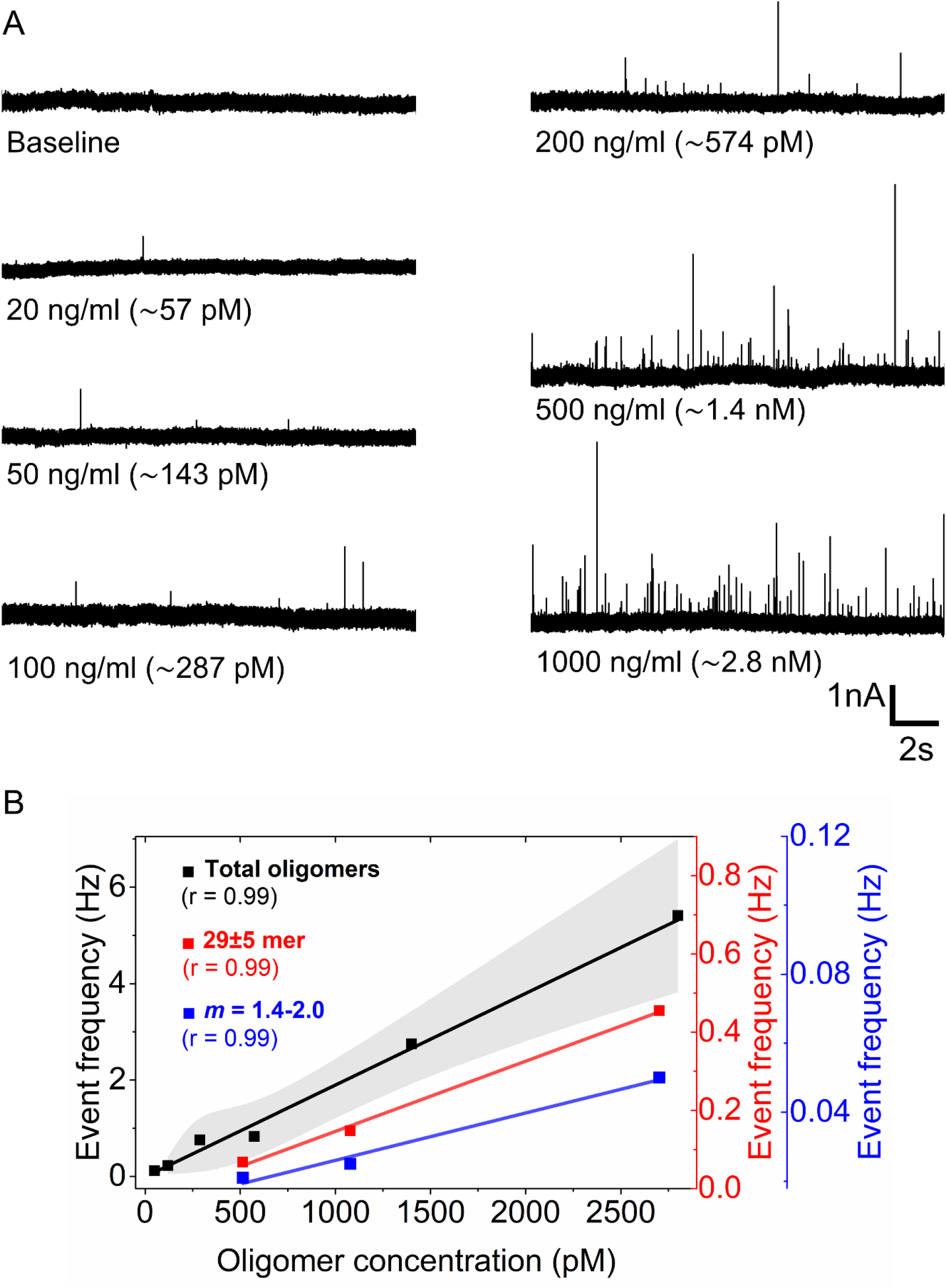
Quantification of the abundance of αSyn oligomers using nanopores. **A.** Original current recordings (~20 s) low-pass filtered with a cut-off frequency of 50 kHz showing resistive pulses (upward spikes) due to translocation of individual αSyn oligomers at increasing total concentrations ranging from 57 pM to 2.8 nM. **B**. The frequency of translocation events at different total oligomer concentrations (black) and frequency of oligomer sub-populations with specific size (29±5-monomers, red) or specific shape (length-to-diameter ratio, *m* ranging from 1.4-2.0, blue). Translocation event frequency is plotted as squares (black, red and blue) and the lines with corresponding colors are linear fits (slopes: 2.0E^-3^ (black), 3.5E^-4^ (red), 2.7E^-5^ (blue) and Pearson’s r values: 0.99 (black), 0.99 (red), 0.99 (blue). The shaded region (light red) shows the standard deviation of four different measurements using a different nanopore chip for each measurement.

Nanopore-based analysis, therefore, provides proof of principle for characterization and quantification of amyloid oligomers at the single-oligomer level in a mixture that is heterogenous with respect to oligomer size and shape. Measuring unmodified oligomers in solution ensures minimal perturbation of the inherent biophysical properties of these aggregates with the possibility of simultaneous oligomer size and shape analysis of each particle. In addition, the recent development of hydrogel-interfaced nanopores combined with asymmetric salt concentrations may enable quantification of particle concentrations in the femtomolar range,^66^ an enabling characteristic for biomarker analysis.

## Conclusions

The work presented here demonstrates that nanopore-based characterization of αSyn oligomers can overcome some of the current challenges in oligomer characterization using conventional methods by providing (i) analysis of the size and shape of oligomers simultaneously, (ii) characterization of oligomers on a single-particle level, thereby circumventing the complications from averaging heterogenous samples, (iii) measurements in solution and in a label-free manner, thereby examining oligomers in their native hydrated state, (iv) quantification of concentrations of total oligomer content or of specific sub-populations in size and or shape that may be particularly toxic or diagnostically relevant. Since oligomers are considered to represent the key neurotoxic species,^8, 9, 20^ a quantitative understanding of their abundance, physical property (i.e., size and shape) and heterogeneity in solution is essential to elucidate their structure-toxicity relationship. The nanopore-based approach presented here enables single-oligomer level, real-time size and shape estimation with minimal sample perturbation, in a heterogenous mixture and it characterizes oligomer samples within tens of minutes based on hundreds of translocation events; even at a total oligomer concentration as low as 150 pM more than 100 translocation events are recorded within 10 minutes.

Determination of size-distributions by the nanopore-based approach revealed 10 stable sub-populations of oligomers as well as approximations of their shape on a single-particle level. This size resolution is twice that of mass photometry and TEM analysis. In addition, TEM-based characterization of oligomers and its potential use for biomarker analysis is typically not available in clinical settings, is expensive and may involve artifact-prone dry-state sample preparation. Importantly these established single-particle methods do not have the capability to determine the shape of individual oligomeric species. Although the nanopore-based approach presented here currently requires specialized training and expertise, it provides rapid, quantitative, label-free, and cost-effective multiparametric characterization of heterogeneous amyloid oligomers in solution. These benefits address the urgent need for methods that make it possible to quantify amyloid oligomers as biomarkers or for testing drug targets to treat neurodegenerative disorders.^32, 36, 67^

## Experimental

### Materials

Oligomers of α-Synuclein were obtained from ND BioSciences, Switzerland as a part of a collaboration supported by the Michael J. Fox Foundation (MJFF-009813) MJFF. The preparation of αSyn oligomers has been discussed in detail by Kumar et al.^55^ Oligomers of αSyn from ND Biosciences were aliquoted, flash frozen, and stored at the −80° C upon arrival and only taken out before the measurements. We obtained the EM-Tec Formvar Carbon support film on copper 200 square mesh (Cat. no. 22-1MFC20-100) for TEM imaging of αSyn oligomers from the company Micro to Nano, Netherlands. All the nanopores used in this work were purchased (pre-fabricated by Helium Ion Milling) from Norcada Inc., Canada or fabricated by helium ion beam sculpting, as explained previously at the University of Arkansas, USA.^68^ The pre-fabricated nanopore chips (4×4 mm) obtained from Norcada contained 10×10 μm SiN_x_ window in a 30 nm thick free-standing SiN_x_ membrane. The chips contain a 100 nm thick SiO_2_ underlayer to reduce capacitive noise. We purchased all other chemicals and buffers from Sigma-Aldrich unless otherwise stated.

### Resistive pulse sensing using nanopores

We carried out all resistive pulse sensing experiments using a recording buffer containing 2 M KCl, and 10 mM HEPES buffer (pH 7.4) as described earlier by Yusko et al.^50^ and Houghtaling et al.^51^ We used polymer-coated solid-state nanopores of 25 nm to 56 nm diameter to characterize αSyn oligomers in free translocation mode as described earlier.^51^ We applied a constant potential of ±100 mV across the nanopore and then measured the current at a sampling rate of 500 kHz via USB-6361 using an AxoPatch 200B patch-clamp amplifier (Molecular Devices) in voltage-clamp mode (β = 1) in combination with LabVIEW (National Instruments) software. We filtered the acquired data with a Gaussian low-pass filter at a cut-off frequency of 50 kHz. We performed a threshold-search (5× the standard deviation of the baseline current) for resistive pulses within the current recordings and determined the oligomer size (as volume) and shape (as length-to-diameter ratio, *m*) by so called “intra-event analysis” as described earlier.^50, 51^ For the analysis of shape with the previously described convolution algorithm,^50, 51^ we constrained the spread “σ” the standard deviation of the current trace value to be greater than, or equal to, the standard deviation of baseline noise. We used Ferritin, a perfectly spherical protein, as a standard to identify the best nanopores, which accurately estimate the size and shape of ferritin and could then be used to determine the size and shape of Syn oligomers (**See Figure 1C**). During the size and shape analysis of Syn oligomers, we used nanopores that give a spherical shape value for Ferritin (with *m* value close to 1).

### Transmission electron microscopy

Carbon coated 300-mesh copper grids (Electron Microscopy Sciences, Hatfield) were plasma cleaned for 5 s using an oxygen plasma cleaner (Zepto RIE, Dienner), before pipetting 5 μl of Syn oligomers sample (~100 μg/ml concentration) in PBS buffer, pH 7.4 on top of the grids followed by 2 min incubation. The grids were washed in a water droplet. Uranyl acetate (2% w/v) was added (3 μl) and incubated for 2 min. Excess stain was blotted off with a filter paper and dried. TEM images were recorded with a FEI Tecnai Spirit operating at 120 kV. We used ImageJ for particle size analysis of Syn oligomers.^69^

### Mass Photometry

We acquired all the mass photometry data using a TwoMP mass photometer (Refeyn Ltd, Oxford, UK) similar to that described previously by Sonn-Segev et al. with slight modifications.^58^ Briefly, microscope coverslips (24 × 50 mm Thorlabs, Cat. No. CG15KH1) cleaned using isopropanol followed by pure water thrice sequentially were used along with PDMS CultureWell™ gaskets (Cat. No. GBL103250) for single-oligomer mass analysis. We diluted αSyn oligomers up to a concentration of ~250 ng/ml in PBS or in 2 M KCl immediately prior to mass photometry measurements. To determine the effect of ionic strength, αSyn oligomers were incubated for 90 min or overnight in PBS or 2 M KCl at room temperature. For each mass photometry acquisition, 20 μL of diluted protein was introduced into the flow chamber, and following autofocus stabilization, movies of 180 s duration were recorded. Each sample was measured at least three times independently (n ≥ 3). Mass photometry data acquisition was performed using AcquireMP (Refeyn Ltd, v2.4.1). All mass photometry images were processed and analyzed using DiscoverMP (Refeyn Ltd, v2.5.0).

## Acknowledgements

SA acknowledges financial supported by the Swiss National Science Foundation “SPARK” funding (CRSK-3_195960). MM acknowledges financial support from the Swiss National Science Foundation (Grant number: 200020_197239), Michael J. Fox Foundation for Parkinson’s Research (MJFF-009813) and from the Adolphe Merkle Foundation, Switzerland.

## —Supporting Information—

***Supplementary Note 1:** Monomer volume of α-Synuclein*

We determined the volume of αSyn monomers in solution using polymer-coated solid-state nanopores. **Supplementary Figure S1C** shows the volume estimates of αSyn monomers based on three independent measurements revealing median volumes ranging from 32 nm^3^ to 39 nm^3^. These volume estimates are close to a volume of αSyn monomers of ~30 nm^3^ as determined by SAXS.^S1^ Moreover, we also used the web server 3V^S2^ for assessment of protein volumes using two different three-dimensional structures of αSyn 1XQ8 and 2KKW determined by nuclear magnetic resonance. This approach revealed a volume of ~24 nm^3^. To determine the size of αSyn oligomers, we used the mean αSyn monomer volume of 35 nm^3^ determined from nanopore-based measurements. The size of αSyn oligomers estimated by nanopores ranges from dimers to 150-mers, dimers to 175-mers, or trimers to 218-mers depending on whether we assumed monomer volumes of 35 nm^3^, 30 nm^3^, or 24 nm^3^, respectively for the calculation. The oligomer size range from dimers to 150-mers that we determined based on an αSyn monomer volume of 35 nm^3^ agrees well with previous reports.^S3, S4^

***Supplementary Note 2:** Stability of oligomers during nanopore recordings*

In order to test the stability of αSyn oligomers with respect to their size distribution in the recording conditions of nanopores (i. e. 10 mM HEPES (pH 7.4), 2 M KCl), we incubated oligomers in PBS or in recording buffer (containing 2 M KCl) for 1 h at room temperature. Thereafter, we determined the oligomer size distribution by TEM imaging of both preparations. **Supplementary Figure S2** reveals no significant difference in the oligomer sizes for αSyn oligomers incubated for 1 h in PBS or in 2 M KCl. In addition to TEM analysis, we also used single-particle mass analysis by mass photometry to determine the effect of buffer conditions on the size of oligomers. **Supplementary Figure S3** reveals no significant difference in oligomer mass distribution was evident among oligomer samples incubated for 90 min or overnight in PBS and 2 M KCl at room temperature. Note, although we did not observe significant changes in oligomer size distribution in the case of the αSyn oligomers studied here, recording conditions may need to be optimized for other amyloid-forming proteins.

**Supplementary Figure S1.**
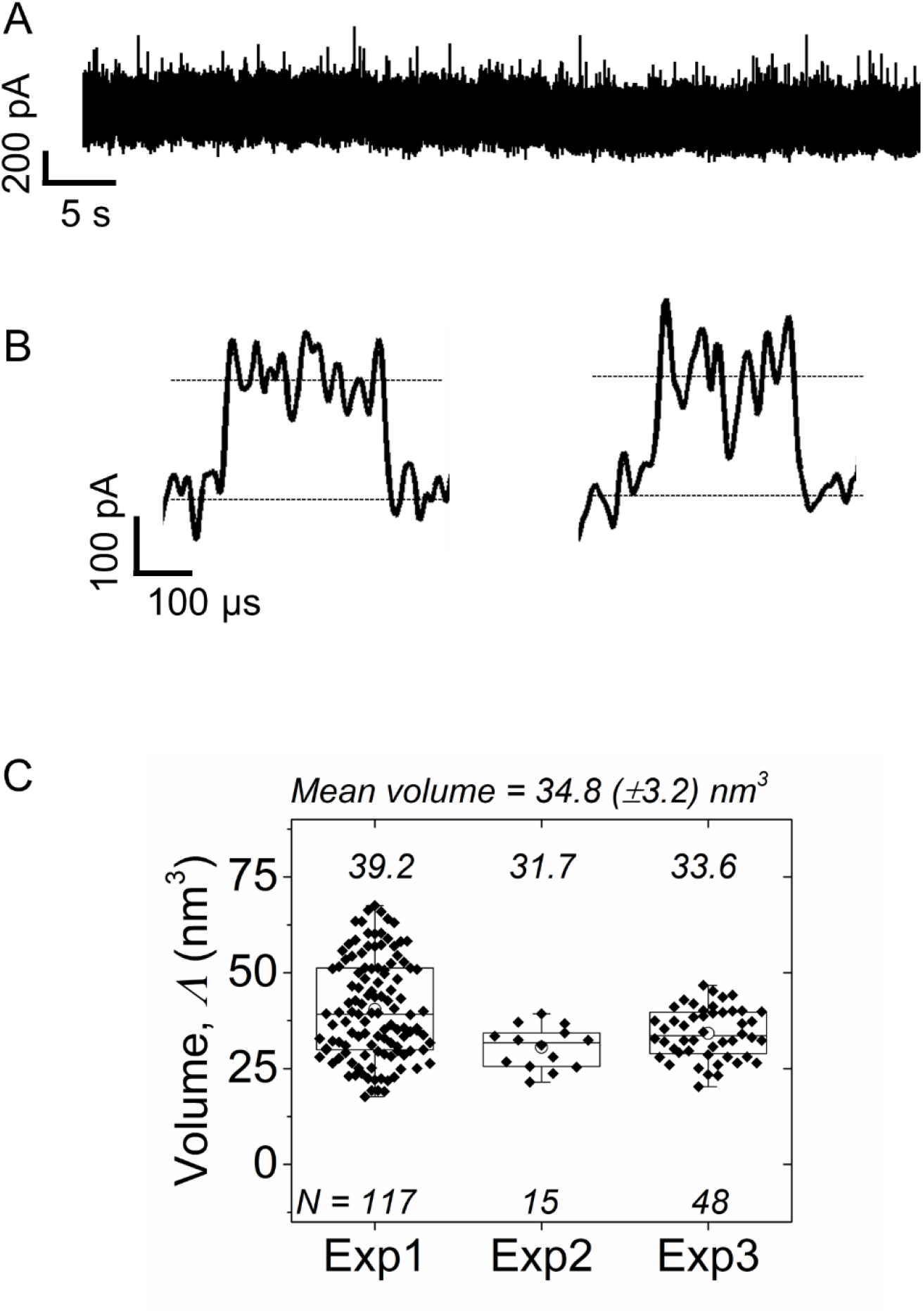
Size of α-Synuclein monomers estimated by resistive pulse sensing. **A.** Original current trace showing individual translocation events of α-synuclein monomers through a nanopore as upward spikes. **B.** Representative individual translocation events of αSyn monomers. **C.** Volume estimates of αSyn monomer from three different nanopore experiments. These results reveal a mean volume of (34.8±3.2) nm^3^. The diamonds show parameter estimates from intra-event analysis with horizontal lines in the box representing the median and quartile values. The mean value is shown as an open circle for each data set, with the whiskers spanning from the 10th to the 90th percentile. The median values for each data set are shown at the top of the corresponding box plot.

**Supplementary Figure S2.**
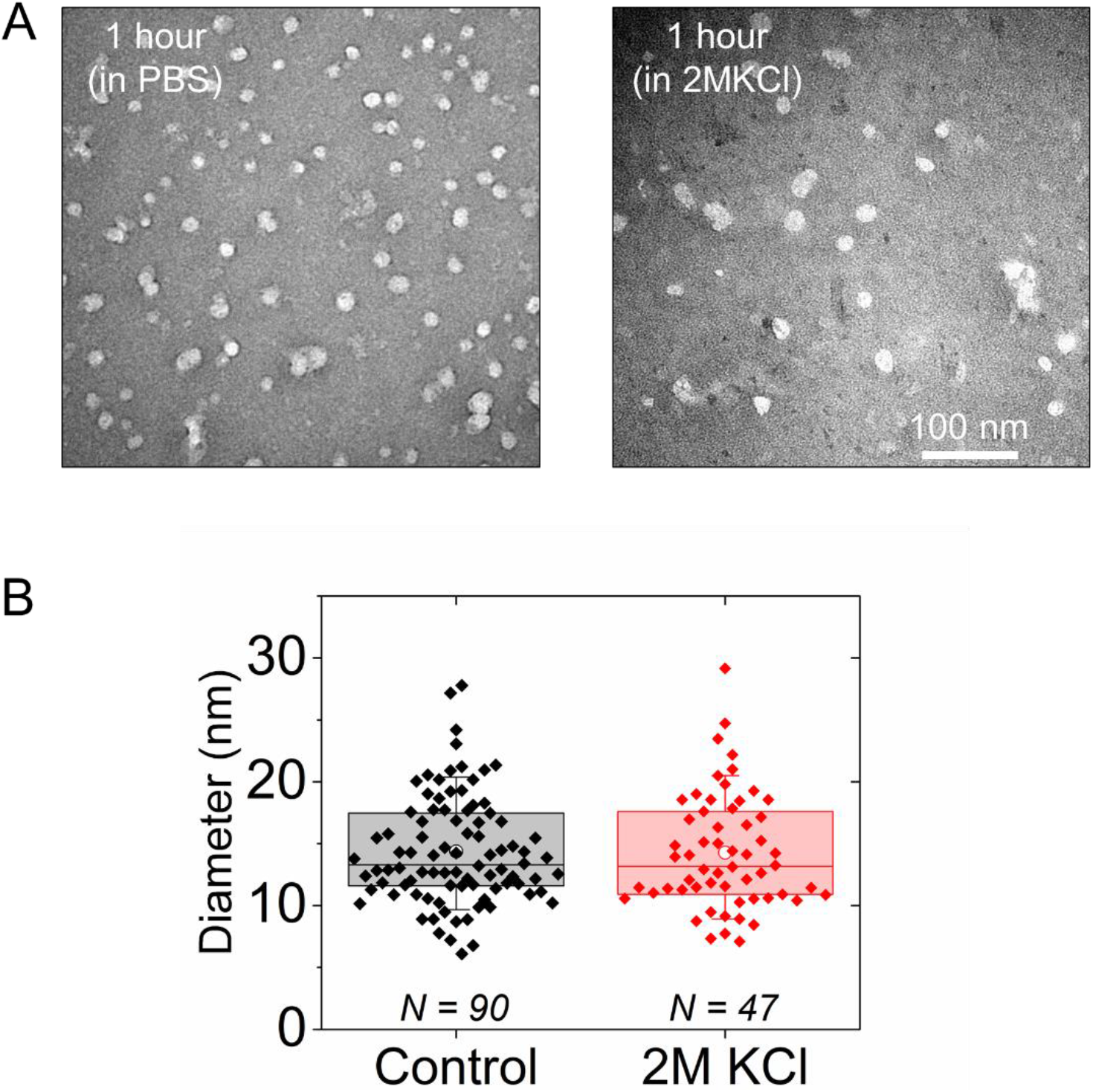
Effect of recording buffer on the size of oligomers determined by TEM imaging. **A.** TEM micrographs of αSynuclein oligomers after 1 h of incubation in PBS (control) or in nanopore recording buffer (i.e. 2 M KCl). **B.** Comparative analysis of oligomer size distribution after incubation in PBS buffer or in 2 M KCl. The diamonds show estimates of oligomer diameter from size analysis of single particles with horizontal lines in the box representing the median and quartile values. The mean value is shown as an open circle for each data set, with the whiskers spanning from the 10th to the 90th percentile. No significant change in oligomer sizes occurred within 1 h of incubation in 2 M KCl.

**Supplementary Figure S3.**
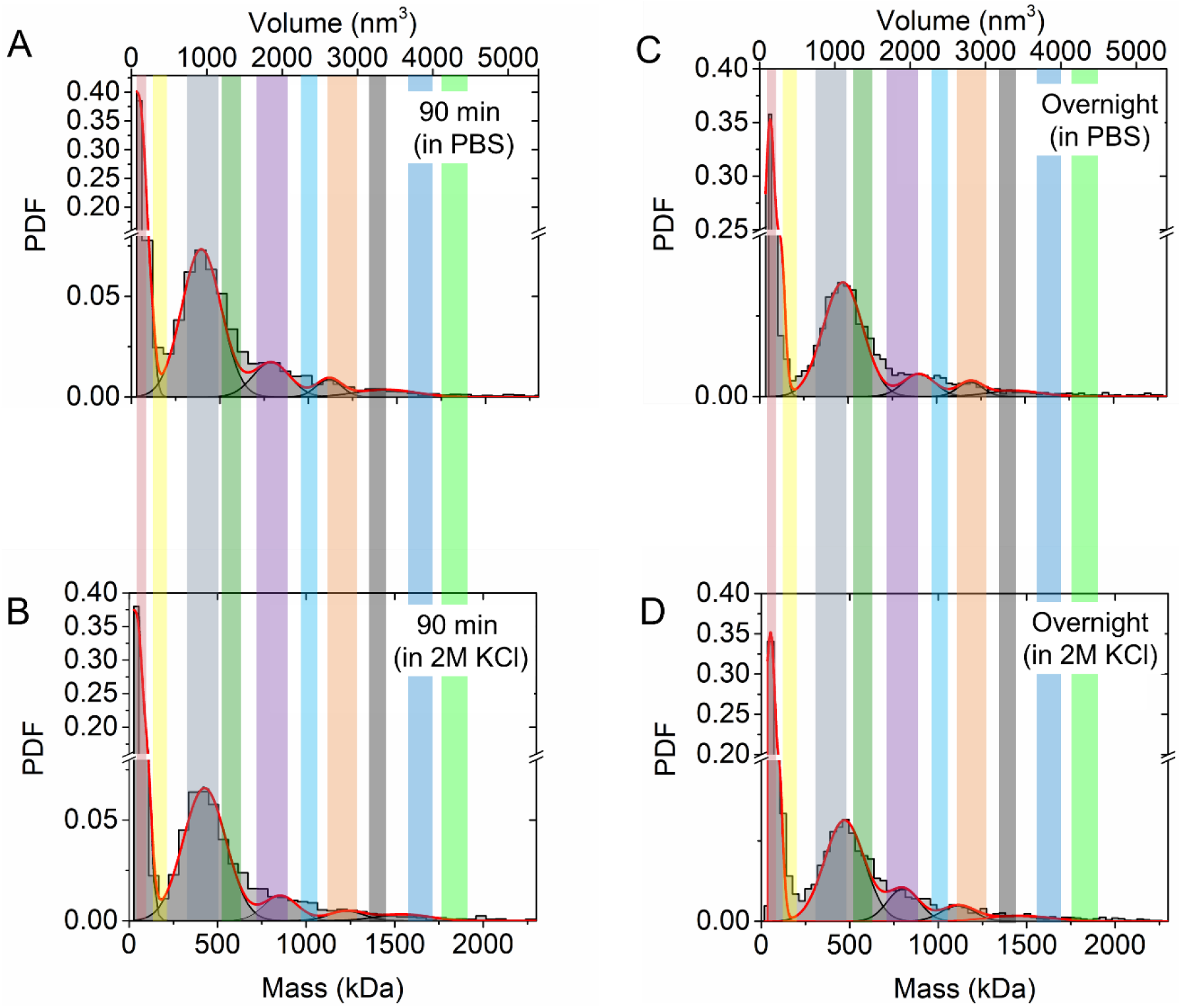
Effect of recording buffer on oligomer size distribution determined by mass photometry. **A.** Size distribution of α-Synuclein oligomers after 90 min incubation in PBS. **B.** Size distribution of α-Synuclein oligomers after 90 min incubation in 2 M KCl. **C.** Size distribution of α-Synuclein oligomers after overnight incubation in PBS. **D.** Size distribution of α-Synuclein oligomers after overnight incubation in 2 M KCl. For comparison, the shaded regions in different colors show the presence/absence of different oligomer populations in different conditions.

